# Integrated Systems Vaccinology Reveals Distinct Metabolic Responses to SARS-CoV-2 Infection and DNA-Based Vaccines in Ferrets

**DOI:** 10.1101/2025.07.11.664150

**Authors:** Gustaf Ahlén, Anoop Ambikan, Flora Mikaeloff, Sefanit Rezene, Jingyi Yan, Negin Nikouyan, Chandrashekar Bangalore Revanna, Sofia Appelberg, Ali Mirazimi, Matti Sallberg, Ujjwal Neogi, Soham Gupta

## Abstract

Understanding systemic effects of vaccination and infection is central to defining correlates of protection against SARS-CoV-2. We used untargeted serum metabolomics to profile the immunometabolic landscape of ferrets after SARS-CoV-2 infection and DNA-/protein-based vaccination. Ferrets were vaccinated with either a multigenic DNA vaccine encoding SARS-CoV-2 RBD, M, and N (OC2), an N-only DNA vaccine (OC12), a recombinant spike protein with QS-21 adjuvant (S+QS21), or a hepatitis B/D control construct (Hep-B/D), and subsequently challenged with SARS-CoV-2. Serum was analyzed longitudinally at baseline, post-vaccination, and post-challenge. SARS-CoV-2 infection induced broad metabolic reprogramming, involving TCA cycle, glutathione metabolism, and nucleotide turnover, reflecting inflammation and cellular activation. OC2 vaccination induced strong metabolic shifts in amino acid and mitochondrial pathways despite low pre-challenge anti-S antibodies. Post-challenge, these shifts extended to redox and nucleotide pathways, correlating with robust anti-S and very strong anti-N antibody responses and complete viral clearance in BAL, but with marked airway pathology, consistent with T cell-mediated clearance of infected cells. S+QS21 and OC12 induced distinct, immunogen-specific signatures with partial protection, while Hep-B/D showed minimal systemic engagement. Metabolite-antibody correlations revealed vaccine-specific associations, highlighting lipid and amino acid pathways as potential immunogenicity biomarkers. Overlap and heatmap analyses showed that metabolic trajectories reflect both the magnitude and quality of immune training. These findings underscore the value of systems vaccinology in resolving mechanistic differences in vaccine responses and support metabolic profiling as a tool for evaluating immune efficacy in preclinical vaccine studies.

## Introduction

The global outbreak of severe acute respiratory syndrome coronavirus 2 (SARS-CoV-2) has resulted in a devastating pandemic with ongoing waves driven by emerging variants of concern^1,2^. Despite widespread vaccination efforts, SARS-CoV-2 remains a major global health threat, particularly in the face of immune evasion and waning protection. In the absence of specific antiviral therapeutics, vaccination continues to be the cornerstone for mitigating disease severity and controlling transmission^3^. Various vaccine platforms such as mRNA, recombinant viral vectors, inactivated viruses, protein subunits, and DNA vaccines have been developed and deployed, with several authorized for emergency use ^4^. Among these, DNA vaccines offer advantages including stability, rapid design, and the potential for inducing robust T cell and antibody responses.

Previously, we demonstrated that DNA-based vaccines encoding the SARS-CoV-2 nucleoprotein (N) and receptor-binding domain (RBD) conferred protection in a ferret model, with coordinated T cell responses and post-challenge antibody production contributing to viral control^5^. Ferrets serve as a well-established preclinical model for SARS-CoV-2, mimicking natural infection in the upper respiratory tract and exhibiting clinical features similar to mild human COVID-19^6^. Furthermore, a recent study in ferrets revealed metabolic alterations particularly in central carbon metabolism during viral shedding ^7^, suggesting that SARS-CoV-2 infection impacts host metabolism in a tissue- and time-dependent manner.

Emerging evidence highlights the importance of metabolism in shaping immune responses to both infection and vaccination. Immune cell activation, clonal expansion, and memory formation are tightly regulated by metabolic programs involving amino acid metabolism, lipid biosynthesis, and mitochondrial remodeling. These immunometabolic shifts not only support effector function but can also leave lasting imprints, contributing to trained immunity or tolerance. As such, the field of systems vaccinology has increasingly leveraged metabolomics to uncover correlates of vaccine efficacy and immunogenicity^8^.

Despite these advances, the systemic metabolic consequences of vaccination and their relationship to immune protection remain incompletely understood, especially in the context of DNA-based SARS-CoV-2 vaccines. Building upon our earlier work in ferrets ^5^, the present study employs untargeted serum metabolomics to characterize the immunometabolic landscape following both SARS-CoV-2 infection and vaccination with four distinct immunogens: a multigenic DNA vaccine encoding RBD, M, and N (OC2), a nucleoprotein-only DNA vaccine (OC12), a recombinant spike protein formulated with QS-21 adjuvant (S+QS21), and a SARS-CoV-2-irrelevant control DNA vaccine targeting hepatitis B and D (Hep-B/D)^9^.

By longitudinally profiling systemic metabolites before and after vaccination, and following SARS-CoV-2 challenge, we identify vaccine-specific metabolic trajectories and correlate them with humoral responses and viral control. This study integrates infection, immunity, and metabolism in a unified framework and provides insights into how DNA-based vaccines modulate host systems biology. Our findings support the application of metabolomic profiling to identify predictive biomarkers of vaccine efficacy and advance the design of next-generation SARS-CoV-2 vaccines.

## Results

### SARS-CoV-2 Infection Induces Distinct Systemic Metabolic Reprogramming in Ferrets

Ferrets are an effective model for studying SARS-CoV-2 infection due to their susceptibility to natural infection, with robust viral replication in the upper respiratory tract and mild disease manifestations resembling those observed in humans^6^. Prior work has demonstrated metabolic shifts in oral, nasal, and rectal swabs of infected ferrets, particularly in central carbon metabolism during peak viral shedding^7^. Building upon this, we investigated the impact of SARS-CoV-2 infection on systemic metabolism using untargeted serum metabolomics.

Six ferrets were used for the study: three were challenged with 10^6^ PFU of SARS-CoV-2 and three with media as mock controls. Baseline serum samples were collected 4 days prior to infection (D0), and post-infection samples (D10) were collected along with nasal wash and bronchoalveolar lavage (BAL) (Figure 1A). Infection was confirmed by RT-PCR targeting the E and RdRp genes in BAL and nasal wash samples, and SARS-CoV-2-specific anti-S and anti-N IgG responses were detected at titers of 1:6250 using an in-house ELISA as described in our earlier study^5^.

**Figure 1.**
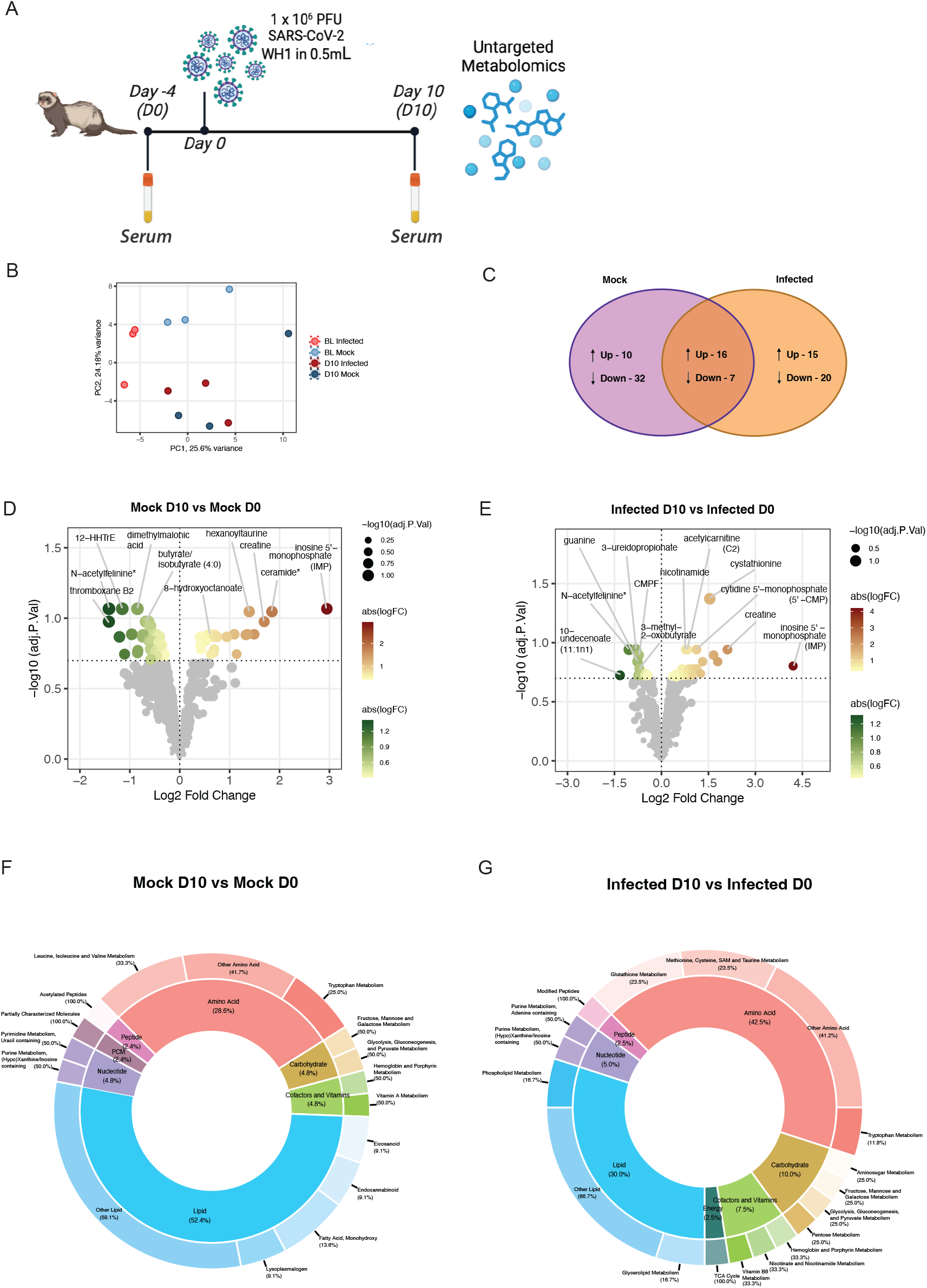
SARS-CoV-2 infection induces systemic metabolic remodelling in ferrets. **A)** Schematic overview of the experimental design. Adult ferrets (n = 3 per group) were intranasally infected with 10^6^ PFU of SARS-CoV-2 or mock-treated. Serum samples were collected at baseline (Day 0, D0) and on Day 10 post-infection (D10). Bronchoalveolar lavage (BAL) and nasal wash samples were collected at D10 for viral RNA quantification, as previously described by Yan et al.^5^. **B)** Principal component analysis (PCA) of untargeted serum metabolomic profiles, illustrating global separation by infection status and timepoint. Each point represents an individual sample; one mock-treated animal showed outlier clustering, likely due to inter-animal variation. **C)** Venn diagram showing the number of significantly altered serum metabolites (FDR-adjusted p < 0.2, LIMMA) between D10 and baseline (D0) in infected and mock-treated animals. Shared and group-specific changes are indicated. **D & E)** Volcano plots showing log_2_ fold change versus -log_10_(adjusted p-value) for metabolites altered at Day 10 relative to baseline in infected (E) and mock-treated (D) animals. Selected metabolites with significant changes (FDR-adjusted p < 0.2) are annotated. **F & G)** Donut plots summarizing the biochemical classification of significantly altered metabolites (Day 10 vs. Day 0) by superpathways (outer ring) and subpathways (inner segments) in infected (G) and mock-treated (F) groups. Percentages indicate the proportion of total significant metabolites belonging to each pathway class.

To investigate the impact of SARS-CoV-2 infection on systemic metabolism, untargeted serum metabolomics was performed using ISO 9001:2015-certified Global Metabolomics (HD4) at Metabolon, NC, USA. Principal component analysis (PCA) revealed moderate separation by both time and infection status, with notable inter-animal variability contributing prominently to sample clustering (Figure 1B). One mock ferret appeared as a strong outlier (rightmost dark-blue), but was retained in subsequent analysis to preserve group size and statistical power.

To correct for inter-individual differences and assess infection-specific metabolic changes, differential expression analysis was performed separately for infected and mock groups, comparing D10 to D0 while adjusting for replicate and time of collection using the LIMMA model. Applying an FDR-adjusted p-value threshold of < 0.2, 51 metabolites were significantly increased and 12 decreased in SARS-CoV-2 infected animals at D10. Among these, 16 upregulated and 7 downregulated metabolites also showed similar changes in the mock group, suggesting these may reflect shared, time-related or procedural effects (Figure 1C, Venn diagram). In contrast, the mock group showed 32 uniquely decreased and 10 uniquely increased metabolites, none of which were altered in infected animals indicating infection-specific metabolic remodeling. These findings, however, must be interpreted cautiously. As seen in Figure 1B, baseline (D0) samples from infected and mock animals were already partially separated, and one mock animal demonstrated outlier behavior. These baseline differences, potentially due to pre-existing metabolic states or individual variation, could influence the observed trajectories. Thus, differential metabolite analyses were carefully adjusted for individual and temporal effects (LIMMA; p < 0.05, FDR < 0.2). The volcano plots showing metabolite changes in both infected and mock groups are presented in Figure 1D.

To further explore the biological context of these changes, differentially abundant metabolites were categorized into superpathways and subpathways (Figure 1E). In mock animals, lipid metabolism (52.4%) and amino acid metabolism (28.6%) dominated, with smaller contributions from carbohydrates, cofactors/vitamins, nucleotides, peptides, and partially characterized molecules. In contrast, infected animals exhibited a marked reduction in lipid-associated changes (30.0%), and a relative increase in amino acid-related (42.5%) and carbohydrate-related (10.0%) metabolites. Additionally, unique infection-driven changes appeared in energy metabolism (2.5%), absent in the mock group. At the subpathway level, SARS-CoV-2 infection enriched metabolic signatures linked to the TCA cycle (100%), glutathione metabolism, methionine-cysteine-SAM-taurine metabolism, and aminosugar metabolism, alongside broader shifts in nucleotide and phospholipid pathways. In contrast, mock animals exhibited subpathway changes involving fatty acids, eicosanoids, and vitamin A metabolism, suggesting these may reflect handling stress or time-dependent metabolic adaptation rather than infection.

### SARS-CoV-2 DNA Vaccines Elicit Distinct Immunometabolic Programs Linked to Antigen Complexity and Viral Control

Ferrets were immunized with plasmid DNA constructs encoding SARS-CoV-2 antigens cloned into the pVAX1 vector, as described previously^5^. The OC2 construct included the receptor-binding domain (RBD), membrane (M), and nucleoprotein (N) of SARS-CoV-2, while OC12 encoded only the N protein. Two control groups were included: a hepatitis B virus/hepatitis D virus DNA vaccine (Hep-B/D) encoding HDAg-P2A-PreS1^9^, used here as a SARS-CoV-2 antigen-negative comparator, and a recombinant spike protein with QS-21 adjuvant (S+QS21), delivered subcutaneously^10^. Ferrets received 300 µg of DNA via intramuscular electroporation using the GeneDrive device (IGEA, Italy), or recombinant spike protein via subcutaneous injection. Serum samples were collected at baseline (T0), Day 31 (T1, pre-challenge), and Day 44 (T2, 11 days post-challenge) with 10^6^ PFU of SARS-CoV-2 (Figure 2A).

**Figure 2.**
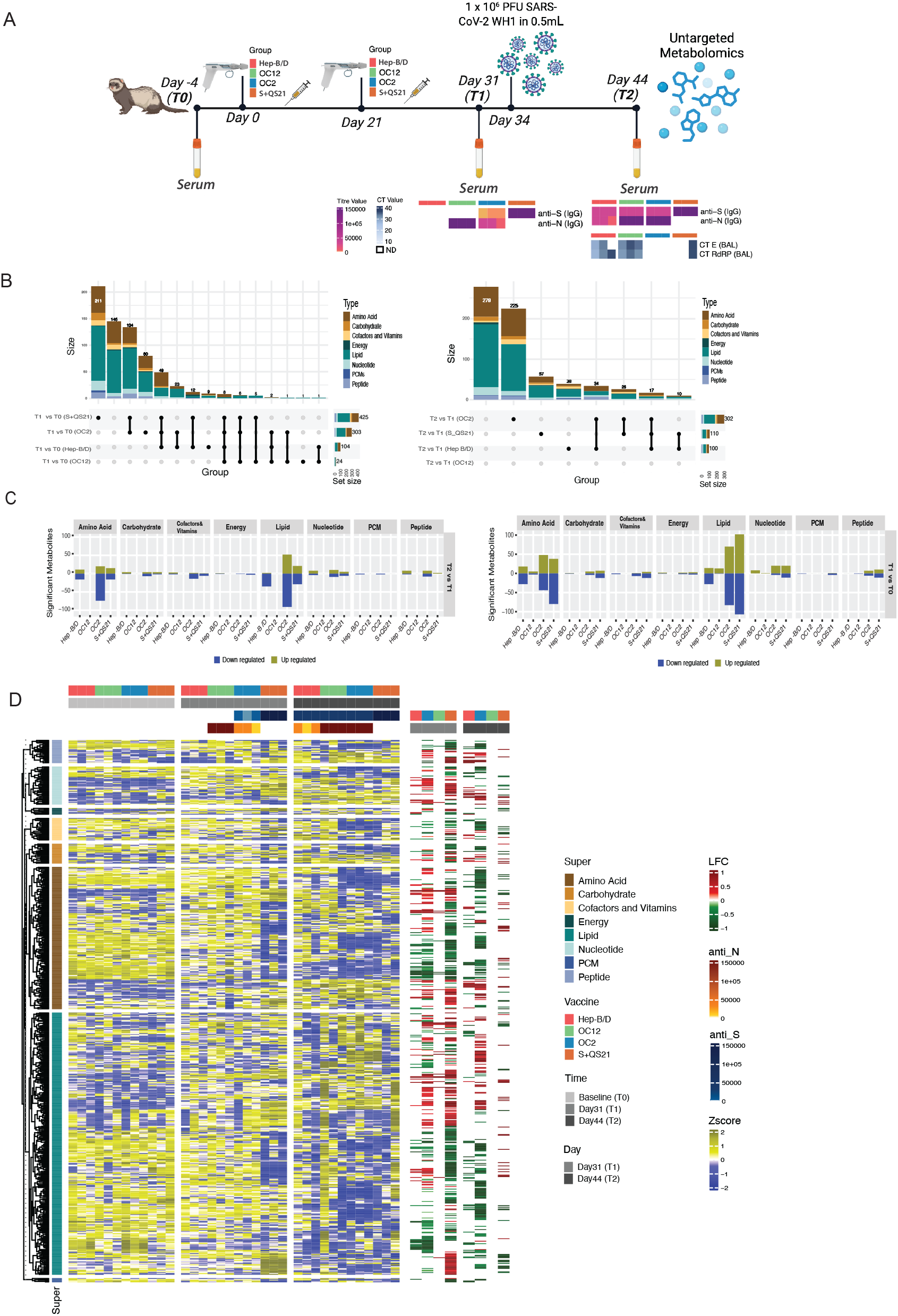
DNA vaccines elicit distinct systemic metabolic responses in ferrets, reflecting immunogen composition and protective efficacy. **A)** Experimental design showing vaccination schedule and sampling timeline. Ferrets (n = 3 per group) were vaccinated with one of four immunogens: multigenic DNA vaccine (OC2), N-only DNA vaccine (OC12), recombinant spike protein with QS-21 adjuvant (S+QS21), or HBV/HDV DNA vaccine (Hep-B/D, antigen-negative control). Serum samples were collected at baseline (T0), Day 31 (T1, pre-challenge), and Day 44 (T2, 11 days post-SARS-CoV-2 challenge). Anti-spike (S) and anti-nucleocapsid (N) IgG titers were quantified by ELISA, and viral RNA in bronchoalveolar lavage (BAL) and nasal wash was measured at T2 by RT-PCR indicated here as heat-map and has been reported earlier by Yan *et al*.^5^. **B)** Bar plots summarizing the number of significantly up- and downregulated serum metabolites by superpathway. The top panel compares T1 vs. T0 (vaccine-induced shifts), and the bottom panel compares T2 vs. T1 (post-challenge responses). **C)** UpSet plots showing overlap of significantly altered metabolites across vaccine groups. Top panel: T1 vs. T0. Bottom panel: T2 vs. T1. Each bar indicates the size of unique or shared metabolite sets across groups. **D)** Heatmap of pathway-level Z-scores for significantly altered metabolites across all vaccine groups and timepoints (T0, T1, T2). Statistical analysis was performed using the LIMMA model, adjusting for repeated measures. Metabolites were considered significantly altered at a Benjamini-Hochberg false discovery rate (FDR) < 0.2. Directionality was determined based on log_2_ fold change.

Principal component analysis (PCA) revealed distinct clustering of vaccinated animals over time. OC2- and OC12-vaccinated groups separated clearly from baseline at Day 31, indicating systemic metabolic remodeling. In contrast, samples from Hep-B/D and S+QS21 groups showed more modest shifts (Figure S1).

Differential metabolomics analysis of Day 31 (T1) samples revealed that OC2 and S+QS21 induced the most extensive changes, with 303 and 425 significantly altered metabolites, respectively (FDR < 0.2), followed by Hep-B/D (104) and OC12 (24) (Figure 2B, left panel upset plot). Although S+QS21 triggered a strong early response, the majority of changes were downregulated, especially in lipid and amino acid metabolism, potentially reflecting acute-phase or inflammatory metabolic suppression. In contrast, OC2’s profile was balanced between up- and downregulated features and enriched in lipid (153), amino acid (92), and nucleotide (20) metabolism (Figure 2C, Left panel metabol count). These metabolomic differences aligned with the breadth of the antibody response. By Day 31, OC2-vaccinated animals had developed strong anti-N IgG titres, while anti-S responses remained low until post-challenge. On the other hand OC12 elicited only anti-N IgG, and S+QS21 only anti-S IgG. Hep-B/D and mock groups remained seronegative at this timepoint (Figure 2A). This dual antibody response in OC2 animals was accompanied by pathway-level activation in amino acid biosynthesis, glutathione and methionine metabolism, and lipid remodeling (Figure 2C, left panel) metabolic programs that support B cell proliferation, antibody production, and redox balance.

By Day 44, following SARS-CoV-2 challenge, OC2 animals continued to display the most pronounced systemic response, with 302 significantly altered metabolites (Figure 2B, right panel, upset plot). This is of particular interest as only the OC2 construct induced T cell responses to RBD, M and N, but did not elicit RBD specific antibodies. These animals also showed the most pronounced histopathological changes in the trachea and carina compared to other groups, and uniquely lacked detectable SARS-CoV-2 RNA in bronchoalveolar lavage (BAL) fluid on Day 10 post-challenge ^5^. Thus, this most likely reflect an effective T cell-mediated clearance of infected cells in the respiratory tissues. This is supported by the dominance of downregulated features, particularly in lipid and amino acid metabolism, is consistent with a transition from immune activation toward resolution and homeostasis. Importantly, after challenge these animals developed and retained strong anti-S and anti-N titres and showed no detectable viral RNA in BAL and reduced levels in nasal wash, confirming effective protection most likely mediated by vaccine induced T cells that eliminated infected cells and promoted rapid antibody responses (Figure 2A). In contrast, OC12-vaccinated animals, despite maintaining high anti-N titres, exhibited no significant metabolic changes post-challenge and had detectable viral RNA in BAL (Ct ∼31-35), indicating incomplete protection. Histopathological analysis also revealed milder inflammation^5^ in these animals, possibly reflecting a less robust T cell response. S+QS21 animals demonstrated moderate metabolic shifts and partial viral clearance (1/3 with detectable virus), while Hep-B/D animals showed low-level, likely non-specific, antibody detection and variable viral burden (Ct 27-36), in line with its role as a negative control.

The specificity of OC2’s systemic response was further highlighted by overlap analysis using UpSet plots, which showed that the majority of altered metabolites were unique to this group (Figure 2C). Shared features between OC2 and OC12 were limited to early lipid changes, and overlap with S+QS21 and Hep-B/D remained minimal (Figure 2C, metabol count, Right panel). Longitudinal assessment of pathway-level changes using Z-score heatmaps reinforced OC2’s distinct metabolic profile. Sustained engagement of pathways related to the TCA cycle, amino acid metabolism, nucleotide biosynthesis, and redox regulation was observed across both timepoints (Figure 2D). These findings suggest a metabolically trained state, supporting continued immune readiness. In comparison, S+QS21 and Hep-B/D exhibited weaker and less coordinated pathway activity, and OC12 largely returned to baseline by Day 44. Metabolite-level insights from volcano plots underscored the breadth and statistical robustness of OC2-induced changes, with large-magnitude shifts spanning multiple metabolic domains. S+QS21 responses were comparatively moderate and scattered. Comprehensive volcano plot analyses of significantly altered serum metabolites (Table S2 and Table S3) are presented in Supplementary Figures S2 and S3. Supplementary Figure S2 shows vaccine-induced changes between baseline (T0) and Day 31 (T1), while Supplementary Figure S3 displays post-challenge changes between Day 31 (T1) and Day 44 (T2) for each vaccine group.

Cumulatively, the OC2 multigenic DNA vaccine induced the most robust and sustained immunometabolic response, characterized by dual antibody specificity, broad metabolic activation, and complete viral clearance. These findings underscore the critical role of antigenic complexity in shaping vaccine-induced systemic immunity. The data further suggest that specific metabolomic signatures particularly those involving amino acid, redox, and lipid metabolism may serve as early biomarkers of vaccine responsiveness and protective efficacy. In contrast, the monovalent N-based OC12 construct and the subunit-based S+QS21 vaccine elicited narrower humoral and metabolic responses, with correspondingly lower viral control. Hep-B/D provided an important baseline for non-specific immune stimulation and confirmed the antigen-specific nature of OC2’s systemic effects.

## Discussion

Our integrative analysis of serum metabolomics in a ferret model of SARS-CoV-2 infection and vaccination reveals immunogen-specific systemic reprogramming, with implications for immune priming, recall responses, and metabolic correlates of vaccine efficacy. These findings build on prior evidence that infection and immune activation are tightly coupled to metabolic rewiring^11,12^, and demonstrate how systemic metabolomics can resolve the qualitative and quantitative nature of host responses across vaccine platforms.

Infection with SARS-CoV-2 induced moderate but targeted metabolic remodelling in ferrets. Notably, infected animals exhibited upregulation of metabolites involved in redox and central carbon metabolism, including cystathionine, creatine, nicotinamide, and inosine monophosphate. These changes were accompanied by pathway-level enrichment in glutathione metabolism, methionine-cysteine metabolism, and the TCA cycle. Together, these shifts suggest activation of oxidative stress buffering mechanisms and mitochondrial energy production, consistent with the metabolic demands of viral replication and innate immune responses^13^. Furthermore, this mirrors earlier observations in COVID-19 patients, where infection perturbs glycolysis, the TCA cycle, glutathione metabolism, and purine biosynthesis ^14-16^. In contrast, mock-infected animals demonstrated broader alterations, particularly in lipid metabolism, including fatty acid and vitamin A pathways. These changes may reflect handling-related stress or time-dependent adaptation rather than specific antiviral responses. Importantly, both groups showed enrichment in tryptophan metabolism, but with higher representation in the mock group and without detection of key downstream products such as kynurenine. This suggests that observed changes in this pathway are likely non-specific and not indicative of interferon-induced immune regulation^17^.

Vaccination elicited substantially more extensive metabolic remodelling than infection alone, with distinct profiles depending on the antigen construct. The multigenic OC2 DNA vaccine (encoding RBD, M, and N proteins) induced broad changes across amino acid, nucleotide, and lipid metabolism before challenge. This metabolic activation was maintained post-challenge and coincided with complete viral clearance and robust dual anti-N and anti-S IgG responses. Enrichment in glutathione, methionine, and amino acid biosynthesis pathways in OC2-vaccinated animals suggests sustained metabolic support for adaptive immunity, including T cell expansion and antibody production^18^. Notably, OC2 vaccination resulted in complete viral clearance and the most dynamic post-challenge metabolome, consistent with its dual antibody specificity and durable immunometabolic imprint.

In contrast, the OC12 vaccine encoding N protein only, although previously shown to be highly immunogenic in murine models^10^, produced limited metabolic shifts pre-challenge and no measurable systemic changes post-challenge, despite eliciting high anti-N IgG titers. These animals retained detectable viral RNA in the lungs, indicating incomplete protection. This dissociation between antibody levels and metabolic activation underscores that humoral responses alone may not reflect or ensure effective immunity^19,20^.

The S+QS21 subunit vaccine triggered the most substantial early metabolic changes, particularly suppressing lipid and amino acid pathways. These effects are consistent with acute-phase responses and the known immunostimulatory properties of saponin-based adjuvants such as QS-21, which can activate dendritic cells^21^ and inflammasomes^22^. However, metabolic responses following viral challenge were less coordinated than in OC2-vaccinated animals, and viral clearance was only partial, indicating that adjuvant-induced inflammation alone may be insufficient to drive sustained protection. One notable observation following viral challenge, with S+QS21-vaccinated as well as OC2-vaccinated animals were enhanced metabolic shifts involving lipid remodeling, redox pathways, and nucleotide metabolism, features associated with immune memory activation and tissue repair. This supports the emerging concept of trained immunity, where prior antigen exposure leads to sustained epigenetic and metabolic programming of innate cells ^23-25^.

Interestingly, the Hep-B/D control group, which received a vaccine formulation devoid of SARS-CoV-2 antigens, exhibited minimal systemic metabolic remodelling. Despite this limited immune engagement, metabolomic analysis revealed an accumulation of purine degradation products and γ-glutamyl amino acids. These metabolites have been consistently associated with chronic inflammation, immune dysregulation, and unfavourable clinical outcomes in COVID-19 patients, suggesting that their presence may reflect a persistent pro-inflammatory state independent of antigen-specific immune activation^26,27^.

Finally, metabolite-level associations with anti-S IgG titers across vaccinated animals suggest that amino acid, redox, and lipid metabolic pathways may serve as correlates of vaccine responsiveness. While causality cannot be inferred, these findings align with recent evidence that metabolic reprogramming in B cells and plasma cells underlies isotype switching and antibody production^28^.

In conclusion, our findings demonstrate that SARS-CoV-2 infection and vaccination drive distinct systemic metabolic programs in ferrets. Whereas infection results in focused metabolic adaptation involving redox and energy balance, multigenic DNA vaccination induces broad and sustained immunometabolic activation linked to effective viral clearance. These results highlight the utility of metabolomics in profiling immune responses and provide mechanistic insights into how antigen composition and vaccine design influence systemic immunity.

## Materials and Methods

### Animal Model, Vaccination, and Challenge Protocol

Ferret vaccination and infection procedures were carried out as previously described in Yan *et al*.^5^. Briefly, eighteen adult ferrets (Marshall Bioresources) were housed under Biosafety Level 3 (BSL-3) containment at the Comparative Medicine Facility (KMF), Karolinska Institutet. Animals were housed in groups of three per cage and acclimatized for 21 days prior to the study. All procedures were approved by the regional animal ethics committee (Ref: 7602-2020) and conducted in accordance with institutional guidelines.

Ferrets were randomly assigned to four vaccine groups. Three groups received DNA vaccines via intramuscular injection (300 µg in the tibialis anterior muscle), followed by in vivo electroporation using the Genedrive device (IGEA, Italy). These groups included: OC2 (encoding SARS-CoV-2 receptor-binding domain [RBD], membrane [M], and nucleoprotein [N]); OC12 (encoding only N); and Hep_B_D (encoding HBV/HDV HDAg-P2A-PreS1, used as an antigen-negative control). The fourth group received recombinant spike protein formulated with QS-21 adjuvant subcutaneously (S+QS21). Vaccinations were administered on Day 0 and Day 21. Two weeks after the final dose, all animals were challenged intranasally with 1 × 10^6^ PFU of SARS-CoV-2 WH1 strain in a 0.5 mL volume. Post-infection, animals were monitored daily for body temperature, clinical signs, and general health.

Serum samples were collected longitudinally at three time points: prior to immunization (T0; baseline), 31 days after the first dose (T1; pre-challenge), and 11 days post-challenge (T2; Day 44). Bronchoalveolar lavage (BAL) and nasal wash samples were collected at necropsy on Day 44 for viral load quantification via qRT-PCR. Additional immunological readouts, including anti-S and anti-N IgG titers, were measured as described previously^5^.

### Untargeted Serum Metabolomics

Untargeted serum metabolomic profiling was performed at Metabolon, Inc. (North Carolina, USA) using the Global Metabolomics HD4 platform, which is ISO 9001:2015 certified. Metabolite extraction and profiling were carried out using ultra-high-performance liquid chromatography coupled with tandem mass spectrometry (UHPLC-MS/MS). Samples were randomized across analytical batches and processed using Metabolon’s proprietary library and software pipeline. Metabolites were identified by comparing chromatographic and mass spectral properties to authenticated reference standards. Only structurally known compounds were included in downstream analyses.

The resultant data matrix included annotated metabolites mapped to superpathways and subpathways based on known biochemical taxonomy.

### Bioinformatic and Statistical Analysis

All computational analyses were performed using R (version 4.4.1) within the RStudio environment. Principal component analysis (PCA) was used to assess global variation between groups and timepoints using the R package PCA tools v2.18.0. Differential expression analysis was conducted using the LIMMA v3.62.2 package^29^, adjusting for repeated measures and within-animal correlations across timepoints. Log2 transformed matrix was used for differential analysis. Comparisons were made between T1 vs. T0 (vaccine effect) and T2 vs. T1 (post-challenge effect). Metabolites were considered significantly altered if they met both nominal p < 0.05 and Benjamini-Hochberg false discovery rate (FDR) < 0.2 thresholds.

Significant metabolites were grouped by superpathway and analyzed for upregulated and downregulated trends. Volcano plots were generated using ggplot2 v3.5.1 package. UpSet plots were constructed using UpSetR v1.4.0 to visualize overlaps of significant metabolites across vaccine groups and timepoints. Heatmaps were created using ComplexHeatmap v2.22.0 to display temporal patterns of pathway-level enrichment.

To explore associations between systemic metabolism and immune readouts, Spearman correlation analysis was performed between normalized metabolite levels and antibody titers (anti-S and anti-N IgG) at both T1 and T2. Correlations were corrected for multiple testing using FDR-adjusted p-values. All plots were generated using base R, ggplot2, or GraphPad Prism 9.

All analyses were conducted blinded to treatment groups to ensure objectivity. Data integration and visualization followed established best practices in metabolomics and systems immunology.

## Supplementary Figures

**Supplementary Figure S1. Principal component analysis (PCA) of serum metabolomic profiles across vaccine groups and timepoints**. PCA plot showing overall variance in serum metabolite profiles across all ferrets, colored by vaccine group (OC2, OC12, S+QS21, Hep-B/D) and timepoint (Baseline, Day 31, Day 44). Principal components 1 and 2 account for 29.9% and 13.2% of total variance, respectively.

**Supplementary Figure S2. Volcano plots of vaccine-**induced **serum metabolite changes between baseline (T0) and Day 31 (T1) across vaccine groups**. Volcano plots showing log_2_ fold change versus -log_10_ adjusted p-value for each vaccine group: **(a)** Hep-B/D, **(b)** OC12, **(c)** OC2, and **(d)** S+QS21. Metabolites significantly altered following vaccination (FDR-adjusted p < 0.2, LIMMA) are displayed, with selected up- and downregulated metabolites annotated.

**Supplementary Figure S3. Volcano plots of post-challenge serum metabolite changes between Day 31 (T1) and Day 44 (T2) across vaccine groups**. Volcano plots showing log_2_ fold change versus -log_10_ adjusted p-value for each vaccine group following SARS-CoV-2 challenge: **(a)** Hep-B/D, **(b)** OC12, **(c)** OC2, and **(d)** S+QS21. Metabolites significantly altered after infection (FDR-adjusted p < 0.2, LIMMA) are shown, with selected annotated features spanning nucleotide, lipid, amino acid, and energy metabolism.

## Supplementary Tables

**Supplementary Table S1:** Differentially abundant serum metabolites in ferrets following SARS-CoV-2 infection.

**Supplementary Table S2:** Differentially abundant serum metabolites in vaccinated ferrets pre-challenge

**Supplementary Table S3:** Differentially abundant serum metabolites in vaccinated ferrets post-SARS-CoV-2 challenge

## Data Availability Statement

All relevant data supporting the findings of this study are included in the manuscript and its supplementary materials. Additional datasets generated and analyzed during the current study are available from the corresponding author upon reasonable request.

## Acknowledgements

This study was supported by the Swedish Research Council through grants 2021-03035 (S.G.), 2021-00993 (U.N.), 2018-06156 (U.N.), and by the OPENCORONA consortium, funded by the European Union’s Horizon 2020 research and innovation programme under grant agreement No. 101003666 (M.S.). M.S. and A.M. acknowledges the support from Swedish Research Council. Additional support was provided by Karolinska Institutet Stiftelser och Fonder.

## Author Contributions

U.N., M.S., S.G. and G.A. conceived and designed the study. G.A. conducted the majority of the experimental work with support from J.Y., N.N., and C.R.B.. A.A. performed the primary bioinformatic and statistical analyses, with contributions from F.M.. S.R. assisted in data interpretation and manuscript preparation. S.A. and A.M. provided expertise in virology and infection models and contributed to study design and sample handling. S.G. and G.A. drafted the manuscript, with additional input from A.A. and S.R. The manuscript was critically reviewed and revised by M.S. and U.N. for important intellectual content. S.G., U.N., and M.S. acquired funding and provided overall project supervision. All authors have read and approved the final version of the manuscript and agree to be accountable for all aspects of the work to ensure its integrity and accuracy. S.G. is the corresponding author and will handle correspondence during the submission and peer review process.

## Declaration of Interests

M.S. is a founder and shareholder of SVF Vaccines AB, which holds patents related to vaccine technologies. G.A. is a consultant to SVF Vaccines AB. All other authors declare no competing interests.

