## Supplementary figures and images for "Integrated Systems Vaccinology Reveals Distinct Metabolic Responses to SARS-CoV-2 Infection and DNA-Based Vaccines in Ferrets"

### Supplemental Figure 1

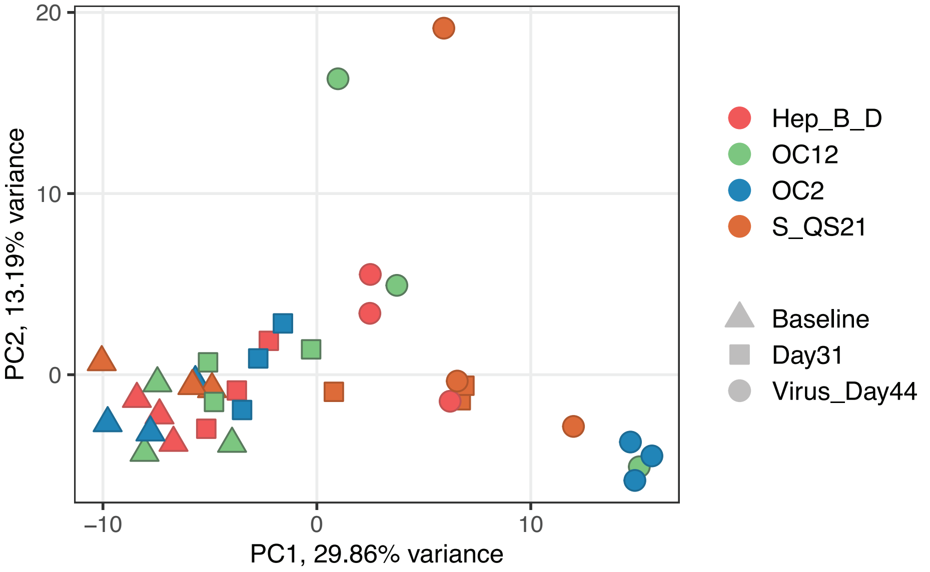

### Supplemental Figure 2

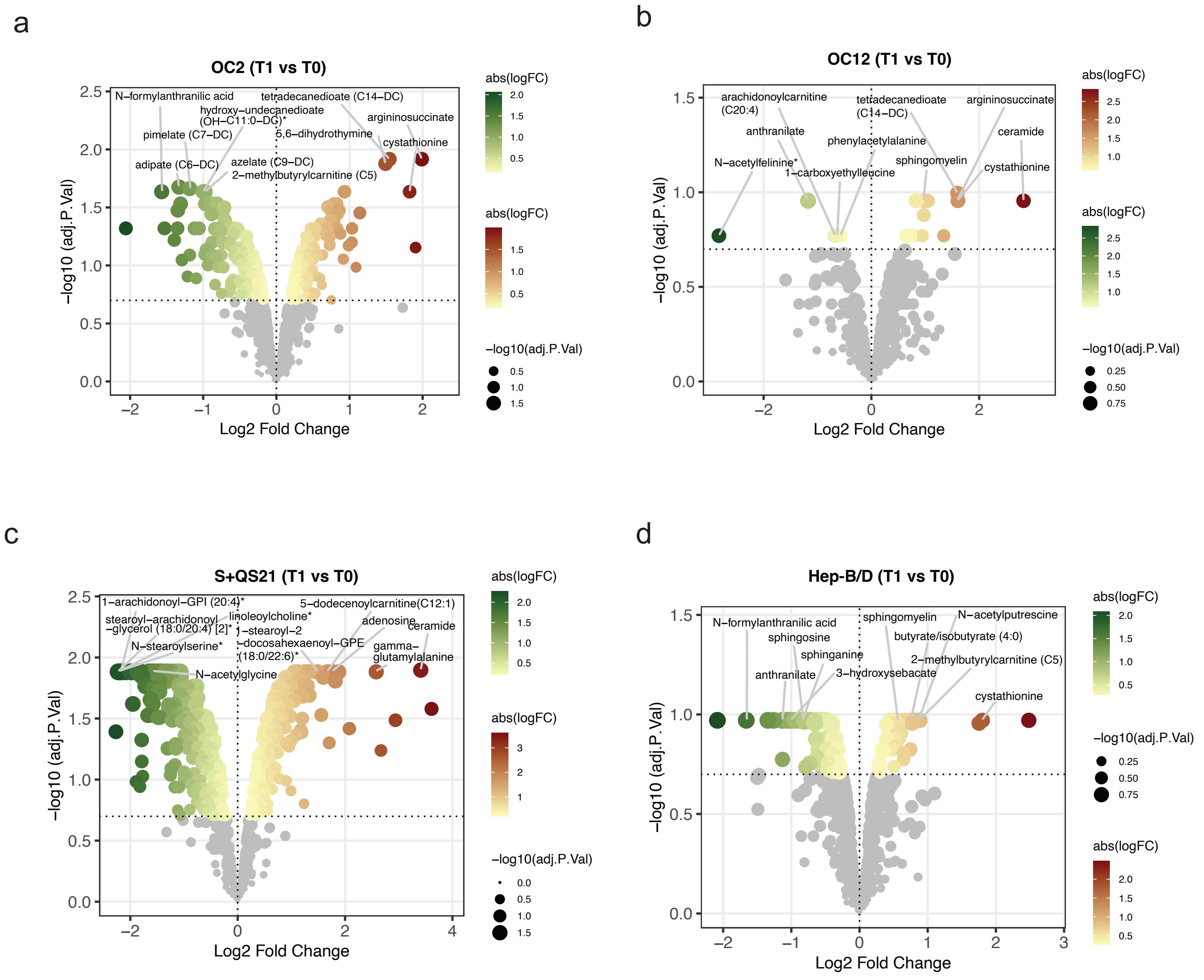

### Supplemental Figure 3

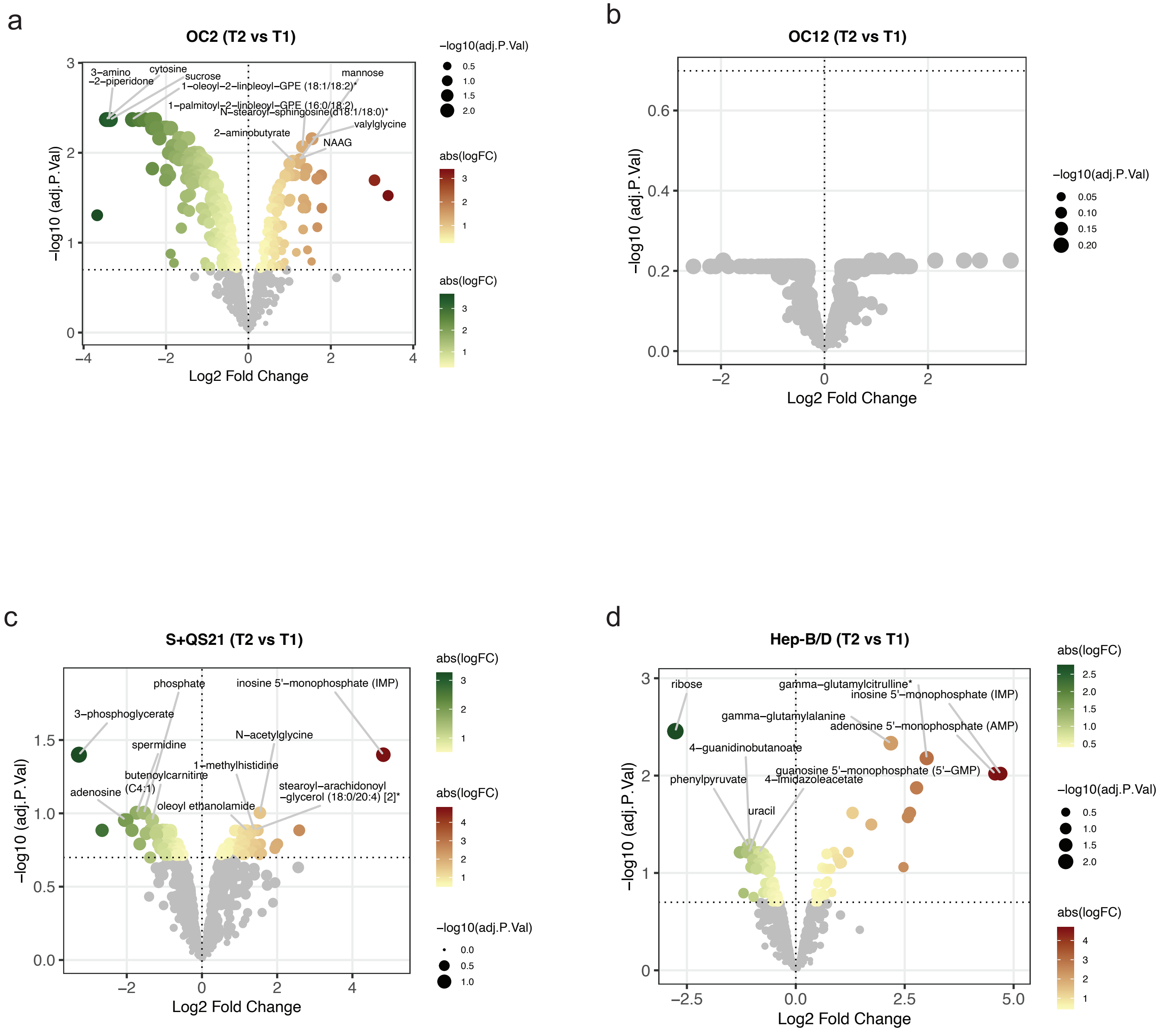
